# Direct microRNA sequencing using Nanopore Induced Phase-Shift Sequencing (NIPSS)

**DOI:** 10.1101/747113

**Authors:** Jinyue Zhang, Shuanghong Yan, Weiming Guo, Yuqin Wang, Yu Wang, Panke Zhang, Hong-Yuan Chen, Shuo Huang

**Affiliations:** State Key Laboratory of Analytical Chemistry for Life Sciences, Nanjing University, 210023, Nanjing, China; Collaborative Innovation Centre of Chemistry for Life Sciences, Nanjing University, 210023, Nanjing, China; School of Chemistry and Chemical Engineering, Nanjing University, 210023, Nanjing, China

**Keywords:** miRNAs, isomiRs, Sequencing, Nanopore, N6-methyladenosine

## Abstract

MicroRNAs (miRNAs) are a class of short non-coding RNAs that function in RNA silencing and post-transcriptional gene regulation. Besides their participation in regulating normal physiological activities, specific miRNA types could act as oncogenes, tumor suppressors or metastasis regulators, which are critical biomarkers for cancer. However, direct characterization of miRNA is challenging due to its unique properties such as its low abundance, sequence similarities and short length. Nanopore Induced Phase Shift Sequencing (NIPSS), which is a variant form of nanopore sequencing, could directly sequence any short analytes including miRNA. In practice, NIPSS clearly discriminates between different identities, isoforms and epigenetic variants of model miRNA sequences. This work demonstrates the first report of direct miRNA sequencing, which serves as a complement to existing miRNA sensing routines by the introduction of single molecule resolution. Future engineering of this technique may assist miRNA based early stage diagnosis or inspire novel cancer therapeutics.

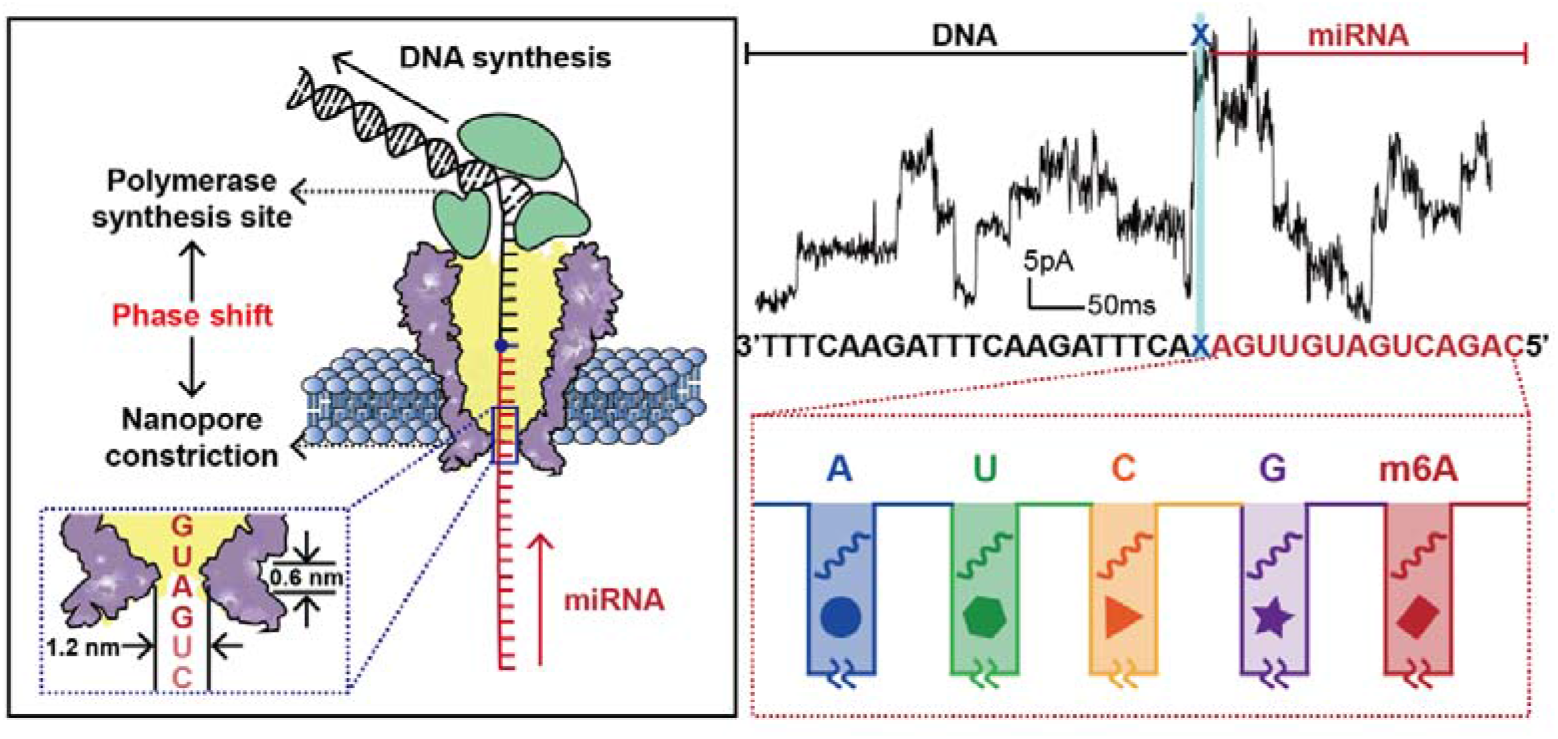

---

MicroRNAs (miRNAs) are a group of short, single-stranded, non-coding RNA molecules that act as posttranscriptional gene regulators for a wide variety of physiological processes, including proliferation, differentiation, apoptosis, and immune reactions^1, 2^. On the other hand, aberrant miRNAs expression levels have been shown to be closely related to diverse diseases, such as cancer^3–6^, auto-immune disorders^7^ and inflammatory diseases^8^.

Conventionally, miRNAs can be characterized by northern blot, quantitative reverse transcription real-time polymerase chain reaction (qRT-PCR) assays or microarrays^9^. Other emerging platforms for miRNA sensing include colorimetry^10^, bioluminescence^11^, enzymatic activity^12^, nanopore sensing^13^ and electrochemistry^14^. Unfortunately, these methods provide limited analytical information because miRNA sequences are not directly reported and prior knowledge of the target miRNA sequence is required.

MiRNAs function by binding to the 3’ untranslated region (3’UTR) of target messenger RNAs^15^. Minor sequence variations, which include trimming, addition or substitution of miRNA sequences, alter its binding affinities to target messenger RNA^15^. As reported, miRNA isoforms (isomiRs)^16^, which were generated by the addition or deletion of one or multiple nucleotides as terminal modifications, have been shown to participate in proliferative diseases^17^ and cancer^18^. On the other hand, N6-methyl-adenosine (m6A), which is an epigenetic modification among RNAs, plays important roles in miRNA biogenesis^19, 20^. These functional roles of miRNAs are associated with their specific sequences. Therefore, directly decoding a miRNA sequence along with its chemical modifications becomes critical to correlate its physiological or pathological relevance. MiRNA could be indirectly sequenced by sequencing its complementary DNAs (cDNA) by performing reverse transcription followed with deep sequencing^21^. However, this strategy suffers from unpredictable amplification biases and the loss of epigenetic information^9, 22^. Advances in corresponding sequencing technologies consequently facilitate early stage diagnosis of cancers and inspire miRNA-targeted therapeutics^23, 24^.

Recent developments in nanopore sequencing have suggested a new concept of nucleic acid sensing by direct sequencing. It has been reported that long stretches of DNA^25, 26^ or RNA^27^ can be directly sequenced by nanopores with single molecule resolution. Unfortunately, miRNAs and other short nucleic acid strands are not compatible with existing nanopore sequencing configurations mainly due to their short length.

Nanopore Induced Phase-Shift Sequencing (NIPSS), which is a variant of nanopore sequencing, was recently developed as a universal strategy to sequence analytes other than long stretches of DNA^25^ or RNA^27^. Though NIPSS is limited by its short read-length of 15 nucleotides, the concept has been verified by sequencing different 2’-deoxy-2’-fluoroarabinonucleic acid (FANA) strands^28^. Mature miRNAs, measuring ∼22 nucleotides (nt) in length, are ideal analytes for NIPSS. Though the 15 nt read-length fails to cover the miRNA length completely, it demonstrates the first single molecule sequencing attempt for miRNAs, which is superior in principle to existing miRNA sensing methods because no amplification or prior knowledge of target sequences is required, while all epigenetic information within the sequence is retained and can be resolved.

## 1. Direct miRNA sequencing using NIPSS

To perform direct miRNA sequencing using NIPSS, the target miRNA strand must be conjugated with a section of DNA to form a DNA-miRNA chimeric template. To prove the feasibility of the method, this chimeric template, which is composed of a segment of DNA on the 3’-end and a segment of miRNA on the 5’-end with the two segments separated by an abasic spacer, was custom synthesized (Fig. 1a, **Table S1**). The sequencing library was constructed by thermal annealing from three separate strands: the chimeric template, the primer and the blocker (Fig. 1a**, Table S1, Methods 1)**^28, 29^.

**Figure 1:**
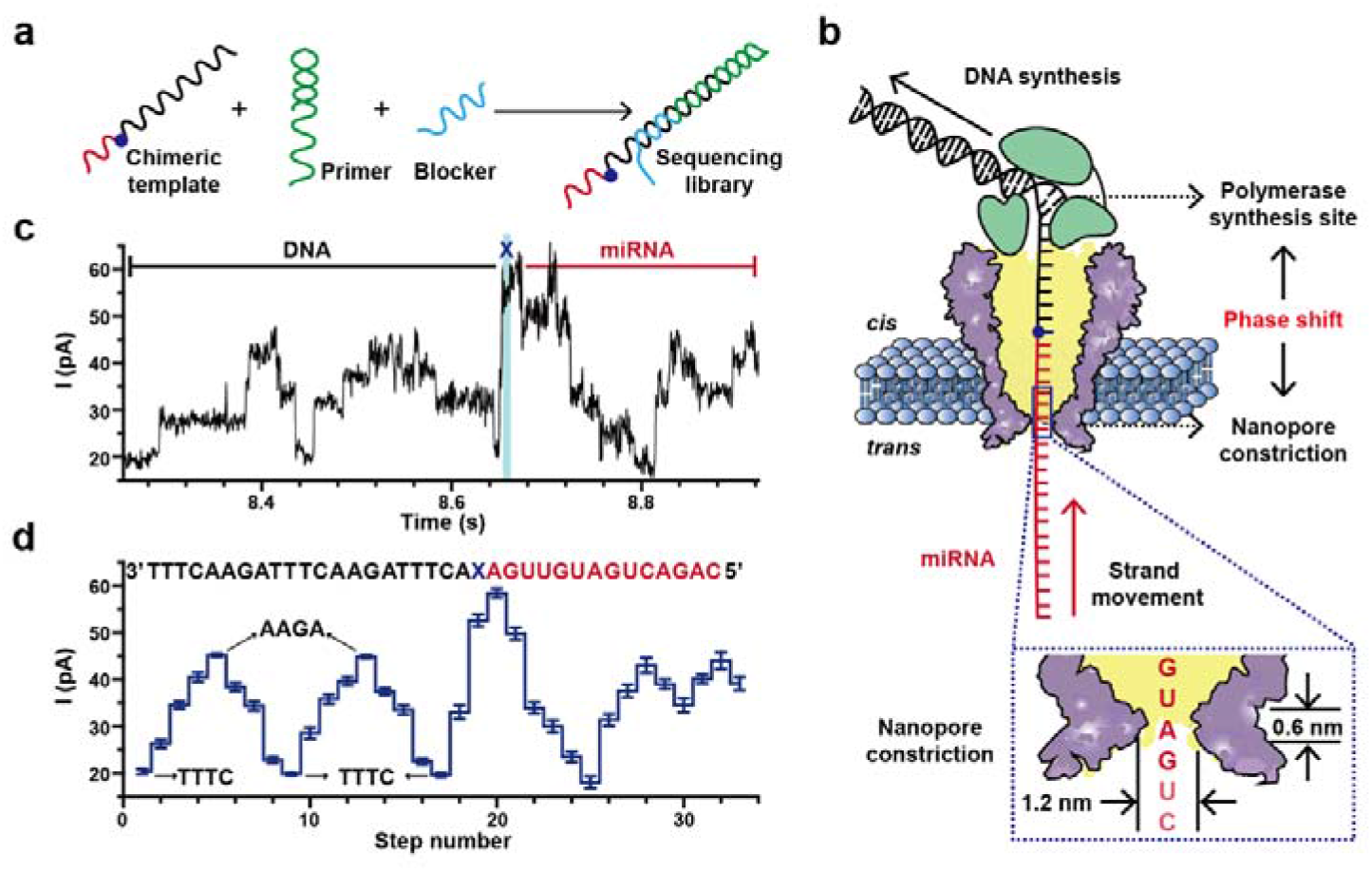
Direct miRNA sequencing using NIPSS. **(a)** A schematic diagram of the preparation of a sequencing library. The miRNA sequencing library is thermally annealed **(Methods 1)** from three separate nucleic acid strands which includes a chimeric template, a primer (green) and a blocker (light blue) **(Table S1)**. The chimeric template is composed of a miRNA segment (red), an abasic residue (blue dot) and a DNA segment (black). **(b)** The NIPSS strategy for direct miRNA sequencing. NIPSS is carried out with an MspA nanopore (purple) and a wildtype (WT) phi29 DNAP (green) by following the reported enzymatic ratcheting strategy **(Fig. S1)**^26^. A fixed phase shift distance between the polymerase synthesis site and the pore constriction is utilized to directly sequence miRNA. During NIPSS, the DNA segment, the abasic residue and the miRNA segment sequentially move through the nanopore constriction in single nucleotide steps until the abasic site (blue dot) reaches the binding pocket of phi29 DNAP. The inset image shows a zoomed-in view of the pore constriction. Due to the limited spatial resolution of the pore constriction, nanopore sequencing signals from MspA results from simultaneous reading of different combinations of sequence quadromers spanning the pore constriction during NIPSS. **(c)** A typical current trace acquired by sequencing DNA-miR-21 **(Table S1)** using NIPSS. The DNA and miRNA segments of the trace are marked with black and red lines, respectively. The trace segment that corresponds to reading the abasic site (X) is marked with a blue stripe. Briefly, the DNA segment of the trace appears as two triangular shaped current characteristics due to the sequence design **(Fig. S2)**. The DNA segment of the trace is immediately followed by an abrupt increase of the signal due to the introduction of the abasic site after the DNA sequence. All step transitions after the abasic signal lead to miRNA sequencing signals. **(d)** Statistics of current steps from sequencing results of DNA-miR-21. The mean and standard deviation values were derived from 24 independent events. The corresponding sequence (3’-5’ convention, if not otherwise stated) is aligned above the statistics. The current levels of “AAGA” and “TTTC”, which represents the highest and the lowest sequencing signals by reading the DNA part, are marked by black arrows. The demonstrated results were acquired by performing NIPSS with an aqueous buffer of 0.3 M KCl, 10 mM HEPES/KOH, 10 mM MgCl_2_, 10 mM (NH_4_)_2_SO_4_ and 4 mM DTT at pH 7.5.

As reported, NIPSS was carried out with a mutant *Mycobacterium smegmatis* porin A (MspA) nanopore **(Methods 2)**, atop of which a wildtype (WT) phi29 DNA polymerase (DNAP) served as the ratcheting enzyme during sequencing^28^. During NIPSS **(Methods 3)**, the sequencing library complex, which is bound with the phi29 DNAP, was first electrophoretically dragged into the nanopore to unzip the blocker strand mechanically. Subsequently, a phi29 DNAP-driven primer extension was initiated so that the chimeric template starts moving against the electrophoretic force in steps equivalent to a single nucleotide **(Fig. S1)**.

As presented in Fig. 1b, the DNA segment, the abasic site and the miRNA segment sequentially pass through the nanopore constriction during NIPSS. The DNA segment thus acts as the “DNA drive strand”. Utilizing the phase-shift between the polymerase synthesis site and the nanopore constriction, the phi29 DNAP enzymatically drives the DNA segment to move against the electrophoretic force and simultaneously the tethered miRNA segment is sequenced by the nanopore constriction. The DNA segment is designed to contain sequence repeats between “AAGA” and “TTTC” (3’-5’) to generate a unique signal pattern during NIPSS **(Fig. S2)**, which marks the initiation of an NIPSS event. The abasic spacer “X” following the DNA segment is expected to produce a high current signature that marks the initiation of miRNA sequencing within a NIPSS event **(Fig. S1, Table S1)**. Since this initiation, miRNA sequencing signals with a read-length of ∼15 nucleotides are expected until the abasic spacer reaches the polymerase synthesis site^28^. Unless otherwise stated, all NIPSS assays described in this paper were carried out by following this configuration. However, the miRNA segments from different chimeric templates contain varying sequences, whereas the DNA segment and the abasic spacer are kept unchanged **(Table S1)**.

As a proof of concept, miR-21, which is an intensively studied oncogenic miRNA and a cancer biomarker^30^, was included in the miRNA segment of a chimeric strand (DNA-miR-21, **Table S1**) for sequencing. A representative raw current trace of DNA-miR-21 is shown in Fig. 1c. This trace can be segmented according to the characteristic sequencing pattern of the DNA drive strand and the abasic spacer respectively, from which the DNA segment reports two repeats of triangular shaped signals and the abasic spacer reports an abnormally high step immediately after the signal from the DNA **(Fig. S2)**. Sequencing signals immediately subsequent from the abasic spacer report sequencing signals for miRNA. Specifically, the representative trace in Fig.1c reports sequencing signals from miR-21.

To analyze the sequencing data **(Methods 4)**, signal steps were extracted from raw sequencing traces by a custom LabView program as reported^28^. The means and standard deviations of all steps were summarized from 24 independent NIPSS events (Fig. 1d). To avoid statistical bias, only NIPSS events with more than 14 nucleotides coverage for the miRNA segment were included **(Methods 4)**. From the statistics, the characteristic high and low current steps which correspond to nanopore readings of AAGA and TTTC were clearly recognized. After the signal from the abasic spacer, 14 subsequent steps that correspond to the additional sequence of miR-21 were demonstrated. Thus, direct miRNA sequencing by NIPSS has been conceptually demonstrated by sequencing a synthetic miR-21 moiety. NIPSS events from synthetic chimeric template containing identical sequences show highly consistent nanopore sequencing patterns, which can be aligned with designed sequences. Direct miRNA sequencing by NIPSS is in principle, universally applicable to other miRNA identities.

## 2. Discrimination of miRNA identities by NIPSS

The sequence of Let-7a, which is a member of let-7 family miRNA^31^, was included to form another chimeric template named DNA-let-7a for sequencing. Let-7 miRNA was the first human miRNA to have been discovered^31^. In contrast to miR-21, which is a cancer biomarker, let-7 miRNA is known to target many oncogenes and thus it behaves as a cancer suppressor^32^. Simultaneous discrimination of miR-21 and Let-7a, which are an oncogenic miRNA and a cancer suppressor miRNA respectively, thus show significant bioanalytical value for cancer diagnosis and serve as an excellent example of miRNA identity recognition by NIPSS.

The NIPSS sequencing of DNA-let-7a was carried out in a manner similar to that demonstrated in Fig. 1. Representative raw sequencing data for DNA-miR-21 and DNA-let-7a are demonstrated in **Fig. S3**. Statistically, an overlay of steps from 24 independent DNA-miR-21 sequencing events (Fig. 2a) and 12 independent DNA-let-7a sequencing events (Fig. 2b) are summarized. The means and standard deviations from the extracted events in Figs. 2a and 2b are shown together in Fig. 2c. The statistics of sequencing steps from both chimeric templates show great alignment in the DNA part of the events. This is expected because the sequence of the DNA segment is identical in DNA-miR-21 and DNA-let-7a **(Table S1)**. Starting from the steps of the abasic spacer however, the sequencing steps appear to deviate from one another due to significant sequence variations in the miRNA segment. Though preliminary, the observed variations of sequencing signals between Let-7a and miR-21 support the hypothesis that different miRNA identifies can be deduced from NIPSS.

**Figure 2:**
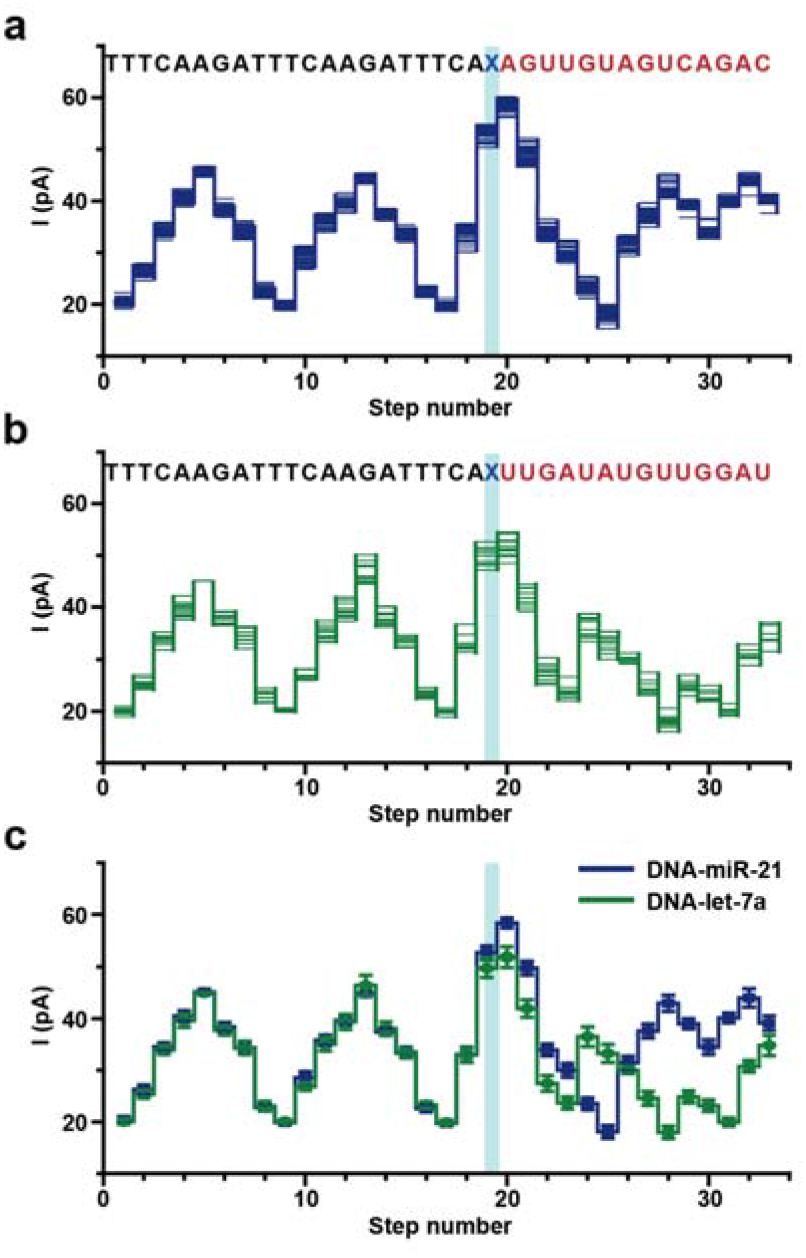
Identification of miRNAs using NIPSS. Chimeric template strands containing miRNA-21 and Let-7a sequences were custom synthesized **(Table S1)** and directly sequenced by NIPSS. **(a)** Overlay of multiple time-normalized events (N=24) from the DNA-miR-21 results acquired by NIPSS. **(b)** Overlay of multiple time-normalized events (N=12) from the DNA-let-7a results acquired by NIPSS. **(a-b)** The corresponding sequences (DNA-miR-21 or DNA-Let-7a, 3’-5’ convention) are aligned above the plots. The DNA segments are marked in black and the miRNA segments are marked in red. The abasic site (X, blue), which separates the DNA and miRNA segments, acts as a signal marker to identify the sequence transition from reading DNA to miRNA during NIPSS. **(c)** Consensus sequencing results comparison between DNA-miR-21 and DNA-let-7a. The mean and standard deviation values are derived from time-normalized events, as demonstrated in **a** and **b**. The DNA part of the NIPSS results shows great alignment in all steps between both templates. However, the miRNA segment of the signals shows significant variations, starting from the step marked with the blue stripe. Blue stripes in **(a-c)** mark the sequencing step of TCAX, which is the first quadromer sequence containing the abasic residue when acquired by NIPSS.

## 3. Discriminating miRNA isoforms by NIPSS

According to the scheme of direct miRNA sequencing by NIPSS, minor sequence variations such as addition, insertion or deletion of nucleotides near the 3’-end of the target miRNA could in principle be resolved by resolution of a single nucleotide. Natural miRNA length isoforms (isomiRs), which result mainly from terminal additions or deletions of an indefinite number of nucleotides at the post-transcriptional level, differ at their 3’ or 5’-end in many mature miRNAs^16^. IsomiRs have attracted significant attention due to their functional roles in diverse biological processes, which include modulation of miRNA stabilization^17^, regulation of mRNA-targeting efficacy^18^ and their correlations to different disease states^9, 17, 18, 33^. Most isomiRs differ by only a single nucleotide at either the 3’ or the 5’-end, and thus discrimination by conventional miRNA sensing routines is challenging. However, isomiRs differing at their 3’-ends as a result of non-template enzymatic additions^34^ are more frequently observed and can be immediately distinguished with the current NIPSS scheme.

The 3’ uridylation product of miR-21, which is an isomiR of miR-21, is named miR-21+U in this paper (Fig. 3a). As reported, natural miR-21+U isomiR is abundant in human urine samples and is a promising biomarker for prostate cancer^35^. As a proof of concept, chimeric template DNA-miR-21 and DNA-miR-21+U **(Table S1)** were designed and sequenced using the same NIPSS configuration (Fig. 1, 2). The means and standard deviations of extracted sequencing steps from 20 independent NIPSS events of DNA-miR-21 and DNA-miR-21+U are shown in Fig. 3b. As expected, the DNA segment of the NIPSS sequencing signal shows perfect alignment, since the DNA drive strands from both chimeric templates are identical in sequence. Distinct variation of the signal begins at step 23 as a consequence of the additional uridine at the 3’-end of miR-21+U when compared to its isoform miR-21. Starting from step 23, all follow-up sequencing steps from miR-21+U appear back shifted by 1 nucleotide (Fig. 3b, c), which further confirms that the 3’ uridylation isomiR has been recognized from direct NIPSS sequencing.

**Figure 3:**
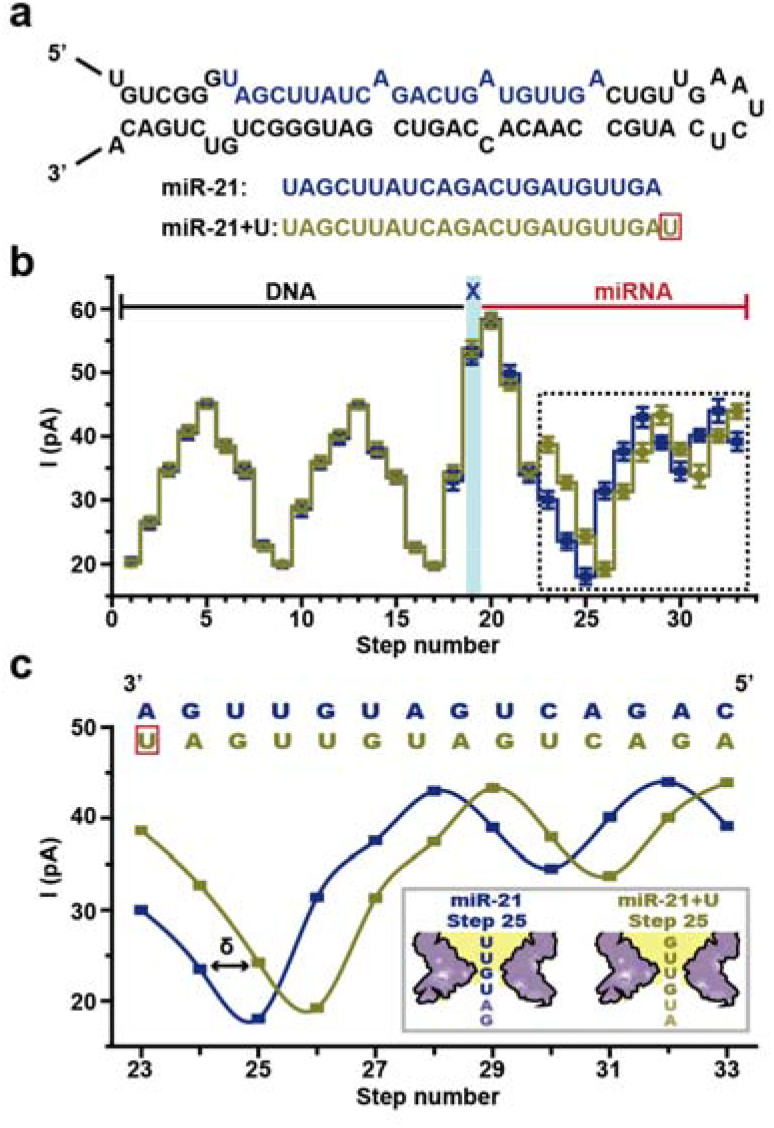
Discrimination of miRNA isoforms (isomiRs) using NIPSS. **(a)** miR-21 and its isoforms. The upper image demonstrates the structure of a precursor microRNA (pre-miRNA) for human miR-21. The lower image demonstrates the sequence (5’-3’) of mature miR-21 and its miR-21+U isoform, respectively. An additional uracil (red box) exists at the 3’-end of miR-21+U. **(b)** Comparison of consensus sequencing results between DNA-miR-21 and DNA-miR-21+U. Consensus sequencing results from NIPSS reading of DNA-miR-21 (brown) and DNA-miR-21+U (blue) derived from 25 time-normalized independent events **(Table S2)**. The DNA, the abasic site and the miRNA part of the signal are marked separately. The blue strip marks the sequencing step of TCAX, which is the first sequence quadromer containing an abasic spacer encountered by NIPSS. The demonstrated statistical results show great alignment in all parts except that marked with a dashed-line box. **(c)** Sequencing result shift between DNA-miR-21 and DNA-miR-21+U. A zoomed-in view of the dashed-line box in **b** illustrates a shift effect of current levels caused by a nucleotide insertion of DNA-miR-21+U in reference to DNA-miR-21. The aligned sequence context above the results demonstrates that the addition of a uracil (red box) in DNA-miR-21+U systematically generates 1 nucleotide (marked as δ) shift. This single nucleotide result shift is also demonstrated by the schematic diagram in the image inset which takes the results of step 25 as an example.

The demonstrated sequencing results between DNA-miR-21 and DNA-miR-21+U have verified that isomiRs with minor sequence variations at the 3’-end can be clearly resolved using NIPSS. As demonstrated, addition or deletion of one or multiple nucleotides can be immediately detected from the characteristic signal variations and the subsequent pattern shift (Fig. 3c). More representative data **(Fig. S4)** and detailed statistics **(Fig. S5)** are shown in the Supporting Information (SI). Minor sequence variations within the first 14 nucleotides to the 3’-end of the target miRNAs can be discriminated by following the same strategy. This makes direct miRNA sequencing by NIPSS immediately applicable to discrimination between members with high sequence similarities from the same miRNA family, such as Let-7^31^.

## 4. Detecting m6A modification within miRNA

Emerging evidence has shown that epigenetic modifications in RNA have a profound influence in gene regulation^36^. N6-methyladenosine (m6A), which is the most abundant modification in RNA, plays a critical role in mRNA metabolism^37^, cellular functions^36, 38, 39^ and miRNA biogenesis^19, 20^. However, due to the chemical and biochemical similarities between adenosine and m6A, precise localization of m6A modifications from natural miRNAs becomes technically challenging for any next generation sequencing platform. The emergence of the technique of MeRIP-seq (methylated RNA immunoprecipitation followed by sequencing)^40, 41^ is a major advance for this technical need. However, MeRIP-seq still requires reverse transcription followed by amplification and fails to achieve single nucleotide resolution. Recent developments of direct RNA sequencing using nanopores have successfully demonstrated m6A mapping directly from mRNA samples^27^. Emerging investigations reveal that m6A modifications also naturally exist in miRNAs^19, 20^. However, direct RNA sequencing by nanopores is only applicable to sequencing long stretches of RNA instead of miRNAs, whereas MeRIP-seq fails to achieve single molecule and single nucleotide resolutions.

NIPSS is particularly useful in distinguishing minor sequence variations at the 3’-end of the target miRNAs. To verify whether single m6A modification within a short stretch of miRNA could be identified by NIPSS, a conceptual experiment was designed. The first nucleotide at the 3’-end of the miRNA segment of DNA-miR-21 is tentatively changed from a canonical adenosine to m6A (Fig. 4a). This new strand is named as DNA-miR-21 (m6A) and is custom synthesized for downstream NIPSS characterization. According to the NIPSS convention, the m6A reaches the pore constriction first (Fig. 4b), which makes this nucleotide highly resolvable by the current NIPSS configuration and thus becomes an optimum option for a proof of concept demonstration.

**Figure 4:**
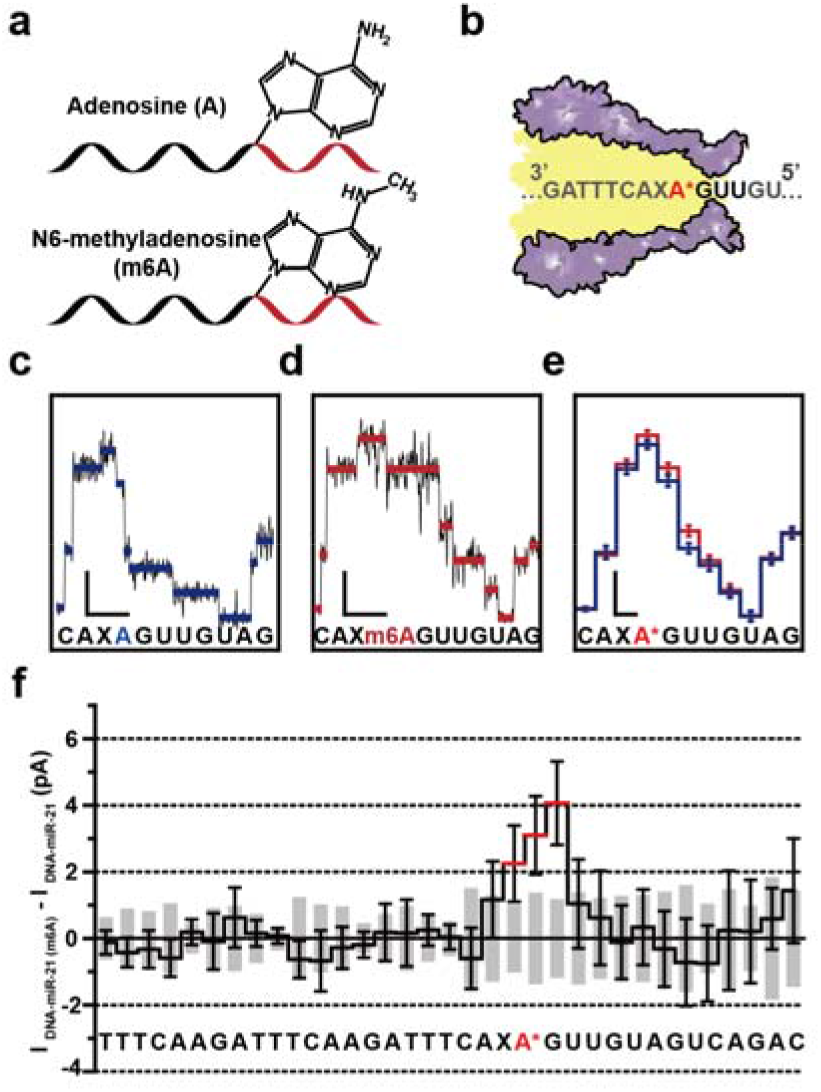
Direct N6-methyladenosine (m6A) mapping using NIPSS. **(a)** Schematic diagram of chimeric template strands containing canonical adenosine (A) or N6-methyladenosine (m6A). A single “A” or “m6A” is embedded in different chimeric strands (top: DNA-miR-21, bottom: DNA-miR-21(m6A), **Table S1**) for NIPSS sequencing, where its chemical structure and location is annotated. Black and red segments represent the DNA and the miRNA part of the strand. **(b)** Schematic diagram of NIPSS sequencing of miRNA containing “A” or “m6A” nucleotides. A* within the sequence context represents the “A” or “m6A” nucleotide within the strand. **(c)** A representative current trace of DNA-miR-21 sequenced by NIPSS. Blue lines represent extracted mean current values from each step. The corresponding sequence context is aligned below in which a canonical adenosine is marked in blue. **(d)** A representative current trace when DNA-miR-21(m6A) is sequenced by NIPSS. Red lines represent extracted mean current values from each step. The corresponding sequence context is aligned below, from which a N6-methyladenosine is marked in red. Scale bars in **c** and **d** represents 10 pA (current, vertical) and 50 ms (time, horizontal), respectively. **(e)** Consensus sequencing results comparison between DNA-miR-21 and DNA-miR-21(m6A). The means and standard deviations were derived from 20 independent events. A* within the sequence context below the results represents either A or m6A. **(f)** Current differences between NIPSS results of DNA-miR-21 and DNA-miR-21(m6A). These differences were derived by calculating Δ*I* = *I*_*DNA−miR*21(*m6A*)_ − *I*_*DNA*−*miR*−21_ from the mean values of 24 events from each strand, with the associated sequence aligned below. The standard deviation values of DNA-miR-21 are demonstrated with gray column and the standard deviation of DNA-miR-21(m6A) is demonstrated with black error bars. The m6A modification results in a signal variation when m6A containing sequence quadromers were read by the nanopore constriction. Due to the limited spatial resolution of MspA, a single m6A modification results in detectable signal fluctuations within three current steps as marked by red lines.

Representative NIPSS sequencing traces from DNA-miR-21 (Fig. 4c) and DNA-miR-21(m6A) (Fig. 4d) show noticeable differences in the current height of the sequencing steps which correspond to the first nucleotide at the 3’-end of the miRNA segment. For a zoomed-in demonstration, only the fraction of the sequencing traces that correspond to the region of interest is shown (Fig. 4c, d) and the additional raw sequencing events are illustrated in **Fig. S6**. The means and standard deviations of the extracted current steps of 24 independent NIPSS events from DNA-miR-21 (blue) and DNA-miR-21(m6A) (red) are summarized and superimposed in Fig. 4e. Due to the limited spatial resolution of the MspA constriction, ∼3 nucleotides show significant signal variations between the two analytes due to the introduced m6A modification, whereas other sequencing steps show a clear alignment as the remainder of the sequence is identical. The difference of sequencing step heights between DNA-miR-21 (m6A) and DNA-miR-21 are shown in Fig. 4f, from which the maximum signal variation between the two analytes could achieve a difference of ∼4 pA in the corresponding positions of the signals, due to the introduced chemical modification **(Fig. S7)**.

## 5. Prospects

The demonstrated miRNA sequencing strategy using NIPSS is currently not without limitations. With the concept of sequencing miRNAs from natural resources by direct miRNA sequencing using NIPSS, the target miRNA strands must be conjugated chemically or biochemically with the DNA drive-strand ahead of the library preparation. An enzymatic ligation strategy has been reported^42^ and could be adapted for this purpose.

Briefly, the 5’ phosphorylated DNA drive strand (5PO_4_ DNA) is first treated with a 5’ DNA adenylation kit (New England Biolabs), assisted with the Mth RNA ligase **(Fig. S8)**. The adenylated DNA drive strand (5AppDNA) was characterized and purified by ethanol precipitation and further ligated to the 3’-end of the target miRNA strands by the T4 RNA ligase 2 truncated K227Q mutant (T4 Rnl2tr). After this ligation, the DNA-miRNA chimeric template can be characterized by electrophoresis on a 15% polyacrylamide gel and the ligated chimeric template is shown as the extra band of higher molecular weight **(Fig. S8)**. In principle, this biochemical conjugation strategy is compatible with sequencing the first 14-15 nucleotides to the 3’-end of any miRNA types. To further extend the read-length of NIPSS to the full length of miRNA, an engineered form of MspA with redundant structures on top of its vestibule could be constructed to extend the phase-shift distance of NIPSS. Alternatively, existing nanopores of larger dimensions such as ClyA^43^ and FraC^44^ may be adapted to sequence full length miRNA or even its precursor or primary form. Chemical ligations ^45, 46^ between the 5’-end of target miRNAs and the 5’-end of the DNA drive strand may be carried out to form a reverse chimeric strand with a “head to head” configuration for 5’-end miRNA sequencing by NIPSS.

## Conclusion

In summary, the first direct miRNA sequencing has been carried out by NIPSS. Similar strategies could also be adapted to sequence other short non-coding strands, such as siRNA^47^ or piRNA^48^, avoiding the laborious motor protein engineering for nanopore sequencing^27^. Though demonstrated as a prototype, the nanopore sequencing results in this paper show clear signal discriminations between different sequences, isoforms and epigenetic modifications among synthetic miRNA sequences. MiRNAs from natural resources, such as miRNA extracts from clinical samples, can be conjugated with pre-designed DNA linker strands by performing routine enzymatic ligation to build NIPSS sequencing libraries **(Fig. S9)**. Consequently, direct miRNA sequencing by NIPSS could be directly implemented in clinical diagnosis or could be utilized as a complement to existing miRNA sensing platforms when single base resolution is critical. MiRNA sequencing by NIPSS shares the same advantages of other nanopore sequencing technologies, including low cost, single base resolution and portability. In principle, NIPSS, when properly engineered could also be adapted to commercial nanopore sequencers, such as MinION^TM^ (Oxford Nanopore Technologies, UK)^27^. Ultimately, high-throughput direct miRNA sequencing by NIPSS can be carried out in optical nanopore chips^49^ for low cost, high-throughput and multiplexed miRNA characterizations in a disposable device form.

## Additional information

Supporting Information (SI). Materials, details of Figures and Tables are summarized in the separate SI PDF.

## Materials & Correspondence

Should be addressed to S.H.

## Acknowledgements

This project was funded by National Natural Science Foundation of China (No. 91753108, No. 21327902, No. 21675083), Fundamental Research Funds for the Central Universities (No. 020514380142, No. 020514380174), State Key Laboratory of Analytical Chemistry for Life Sciences (Grant No. 5431ZZXM1804 Grant No. 5431ZZXM1902), Excellent Research Program of Nanjing University (Grant No. ZYJH004) 1000 Plan Youth Talent Program of China, Programs for high-level entrepreneurial and innovative talents introduction of Jiangsu Province. Technology innovation fund program of Nanjing University.

## Methods

### 1. The construction of miRNA sequencing library

The sequencing library for NIPSS is thermally annealed from three separate nucleic acid strands: the chimeric template, the primer and the blocker (Fig. 1a**, Table S1)**. These three strands were mixed with a 1:1:2 molar ratio in an aqueous buffer (0.3 M KCl, 10 mM HEPES/KOH, 10 mM MgCl_2_, 10 mM (NH_4_)_2_SO_4_). Thermal annealing was carried out by incubating the mixture at 95 °C for 2 min and program cooled down to 25 °C with a rate of −5 °C/min. For optimum sequencing data production, the thermal annealed sequencing library should be immediately used in subsequent NIPSS measurements.

### 2. The preparation of a biological nanopore

The MspA mutant (D90N/D91N/D93N/D118R/D134R/E139K) nanopore^50^ was expressed with *E. coli* BL21 (DE3) and purified with nickel affinity chromatography as described previously^28, 51^. This MspA mutant, which is the sole MspA nanopore discussed in this paper, is named MspA, if not otherwise stated.

### 3. NIPSS experiments

NIPPS experiments were carried out as described previously^28^. Briefly, the electrolyte buffer (0.3 M KCl,10 mM HEPES/KOH, 10 mM MgCl_2_, 10 mM (NH_4_)_2_SO_4_ and 4 mM DTT at pH 7.5) were separated by a 1,2-diphytanoyl-sn-glycero-3-phosphocholine (DphPC) lipid membrane (Avanti Polar Lipids) into *cis* and *trans* compartments. Both compartments were in contact with separate Ag/AgCl electrodes and connected to an Axopatch 200B patch clamp amplifier (Molecular Devices) to form a circuit, while the *cis* compartment is electrically grounded. Purified MspA nanopores were added in *cis* and spontaneously inserted into the membrane. With a single pore inserted (Fig. 1b), the sequencing library, dNTPs and phi29 DNAP could be added into *cis* and stirred magnetically to reach final concentrations of 2 nM, 250 μM and 1 nM, respectively. Nanopore sequencing was initiated by holding an applied voltage at +180 mV. All electrophysiology recordings were acquired with a Digidata 1550B digitizer (Molecular Devices) with a 25 kHz sampling rate and low-pass filtered at 5 kHz. All NIPSS experiments were performed at room temperature (22 ±1 °C).

### 4. Data analysis

All data analysis was performed identically to that in previously reported work using NIPSS^28^. Briefly, trace segments containing NIPSS events were extracted from raw electrophysiology traces. Sequencing steps, which appear as signal plateau transitions within the trace, were extracted by a custom LabView program. The DNA-miRNA chimeric design facilitates further data analysis. The characteristic signal pattern as acquired by sequencing the DNA drive strand and the abasic spacer, clearly marks the initiation of the miRNA sequencing signals afterwards. To avoid statistical biases, all statistics of sequencing results were taken from NIPSS events with more than 14 nucleotides coverage for the miRNA segments.

