## Supporting Information for "Direct microRNA sequencing using Nanopore Induced Phase-Shift Sequencing (NIPSS)"

**Materials**

Potassium chloride (KCl), sodium chloride (NaCl), sodium hydrogen phosphate ($Na_2HPO_4$) and sodium dihydrogen phosphate ($NaH_2PO_4$) were obtained from Aladdin (China). Magnesium chloride ($MgCl_2$) was from Macklin. Ammonium sulfate $(NH_4)_2SO_4$ was from Xilong Scientific. 4-(2-hydroxyethyl)-1-piperazine ethanesulfonic acid (HEPES) was from Shanghai Yuanye Bio-Technology (China). Ammonium persulfate, kanamycin sulfate, dl-dithiothreitol (DTT), dioxane-free isopropyl-β-D-thiogalactopyranoside (IPTG), N,N,N',N'-Tetramethyl-ethylenediamine (TEMED) and imidazole were from Solarbio (China). Ethylene-diaminetetraacetic acid (EDTA), pentane, hexadecane, and Genapol X-80 were from SIGMA-ALDRICH. 1,2-diphytanoyl-sn-glycero-3-phosphocholine (DPhPC) was from Avanti Polar Lipids. Urea was from BIOSHARP. Acrylamide was from Sangon Biotech. The Low Range ssRNA Ladder (#N0364S), microRNA Marker (#N2102S), RNA loading dye (#B0363S), Nuclease-free Water (#B1500S), T4 RNA ligase 2 truncated K227Q mutant (T4 Rnl2tr; #M0242S), phi29 DNA Polymerase (#M0269S), deoxynucleotide (dNTP) solution mix (#N0447S) and the 5' DNA Adenylation kit (E2610S) were from New England Biolabs. *E. coli* strain BL21 (DE3) was from Biomed (China). Luria-Bertani (LB) agar and LB broth were from Hopebio (China).

All HPLC-purified DNA oligonucleotides, including DNA-miRNA template strands, primer, blocker, DNA linker and miR-21+U **(Table S1)** were custom synthesized by Genscript (New Jersey, USA).

The MspA mutant (D90N/D91N/D93N/D118R/D134R/E139K) nanopore was expressed with *E. coli* BL21 (DE3) and purified with nickel affinity chromatography as described previously[1]. This MspA mutant was abbreviated as MspA in the paper.

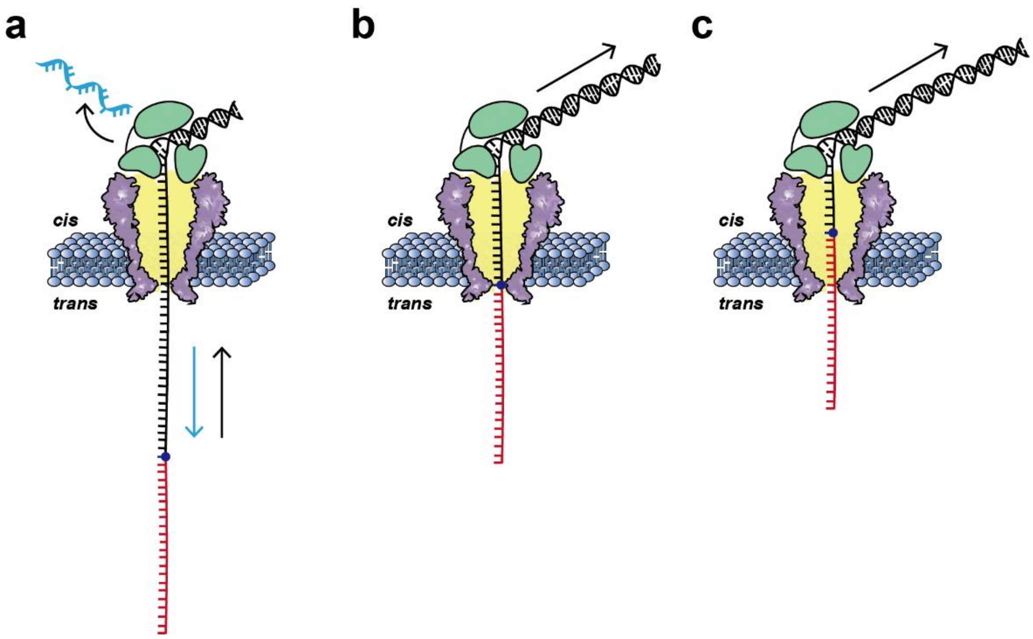

**Fig. S1: Detailed schematic diagrams of NIPSS.** The configuration of direct miRNA sequencing by NIPSS is as reported previously[2]. The chimeric template is composed of a miRNA segment (red), an abasic residue (blue dot) and a DNA segment (black) **(Methods 1)**. With a +180 mV applied potential, the sequencing library, which was bound with a phi29 DNAP (green), was driven electrophoretically into the MspA nanopore (purple). **(a)** Initiation of NIPSS. The cyan DNA blocker strand, which was thermally annealed with the sequencing library, was first voltage-driven fragment unzipped from the chimeric template. This unzipping consequently triggered the replication-driven ratcheting by the phi29 DNA polymerase on top of the nanopore. The transition from voltage-driven unzipping to replication-driven ratcheting represents the initiation of nanopore sequencing. The motion directions of the chimeric template during unzipping and ratcheting are marked with cyan and black arrows, respectively. **(b)** The initiation of miRNA sequencing. Passage of the abasic spacer through the pore constriction marks the initiation of subsequent miRNA sequencing. **(c)** MiRNA sequencing by NIPSS. By utilizing the phase-shift, the miRNA segment passes through the pore constriction with single nucleotide steps, when the DNA drive-strand is being replicated by the phi29 DNA polymerase. All NIPSS experiments in this paper follow this configuration. All sequencing experiments were performed at 23 $^{o}C$ in the sequencing buffer of 0.3 M KCl, 10 mM HEPES/KOH, 10 mM $MgCl_2$, 10 mM $(NH_4)_2SO_4$ and 4 mM DTT at pH 7.5.

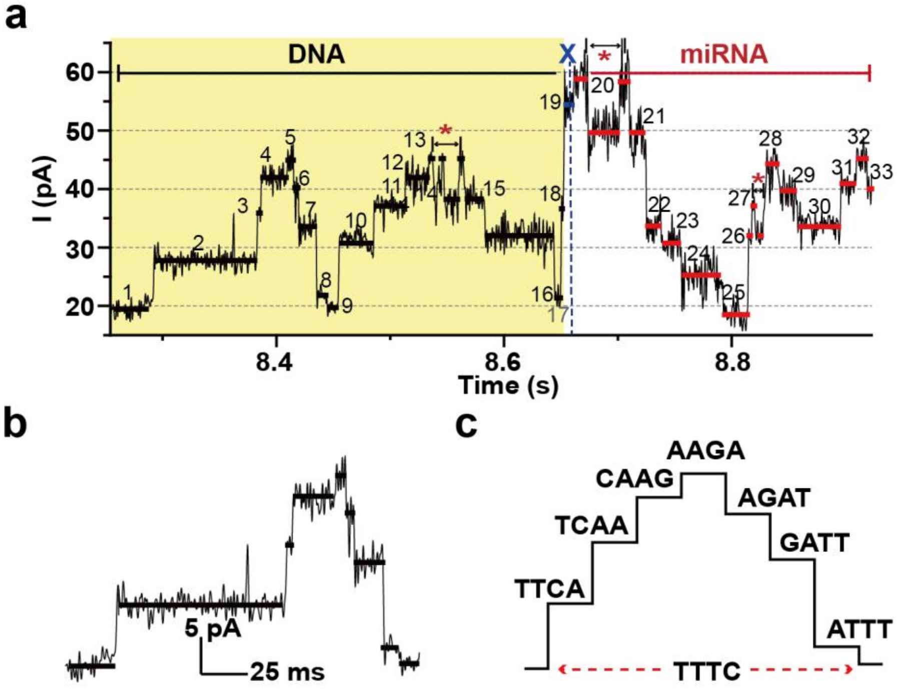

**Fig. S2: Design of the DNA segment within the chimeric template. (a)** A representative current trace acquired by sequencing DNA-miR-21 using NIPSS. The trace segments that correspond to reading DNA, the abasic spacer and miRNA are marked appropriately. The purpose of including the DNA segment, which is designed on the 5'-end of the chimeric template strand **(Table S1)**, is twofold. First, it acts as a drive strand, which can be enzymatically ratcheted by the phi29 DNAP against the electrophoretic force, guarantees that the miRNA segment can be sequenced by nanopores. Second, by including two sequence repeats of "AGAACTTT" (5'-3') in the DNA segment **(Table S1)**, a unique trace pattern of two triangular wave (region marked with yellow shade) will appear ahead of the miRNA reading, which helps to recognize the initiation of an NIPSS event. The assigned numbers above each trace plateaus represent different quadromer readings during NIPSS. Nanopore reading of AAGA and TTTC (3'-5') generates the highest and the lowest residual current among that from all other canonical combinations of DNA quadromers respectively. Thus, quadromer reading from AAGA could be recognized as steps 5 and 13. Whereas, quadomer reading from TTTC could be recognized as steps 1, 9 and 17. Back-stepping motion[3] of the nucleic acid strand, was occasionally observed, and marked with "*". **(b)** The characteristic current signature generated by the DNA segment. The demonstrated trace was extracted from step 1-9 from **(a)**. Scale bar: 5 pA/25 ms. **(c)** The extracted current steps from **(b)** with aligned quadromer sequences. The 14 sequencing steps, which corresponds to steps marked between 20-33, contain miRNA sequencing signals.

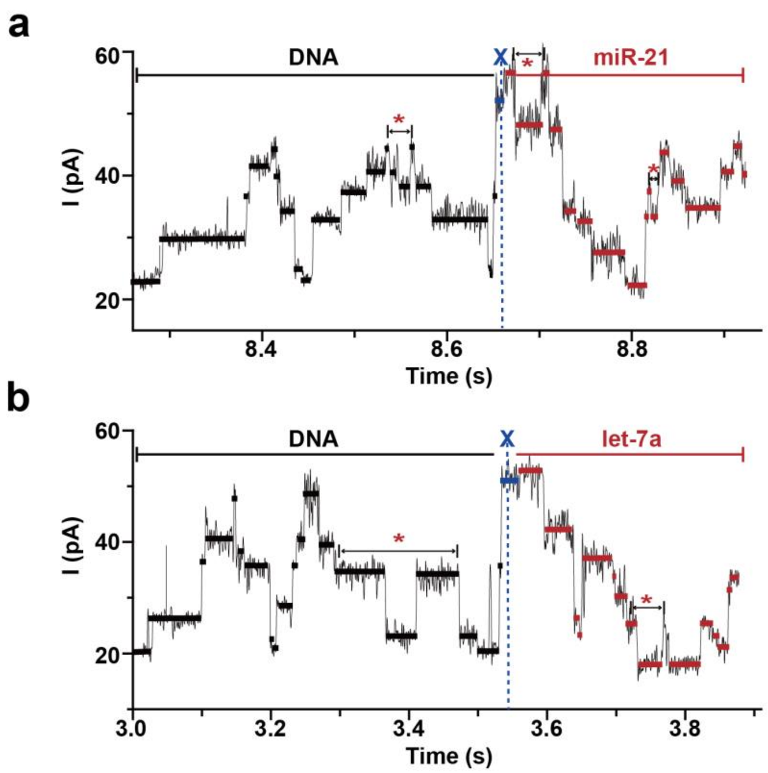

**Fig. S3: Identification of miR-21 and let-7a by NIPSS.** MiRNA identities can be directly recognized by analyzing the miRNA part of the NIPSS signal. **(a)** A representative current trace from DNA-miR-21 sequencing. **(b)** A representative current trace from DNA-let-7a sequencing. Solid lines over the traces in **(a-b)** represent the extracted current height. "*" indicates occasional polymerase back-stepping[3]. The initiation of miRNA sequencing is indicated by blue dashed lines, where an abasic spacer (X) is read by the nanopore. The signal patterns of the DNA segment show high similarities since the sequence of the DNA drive-strand is identical. Whereas, the miRNA segment of the signal show remarkable differences between the demonstrated NIPSS events.

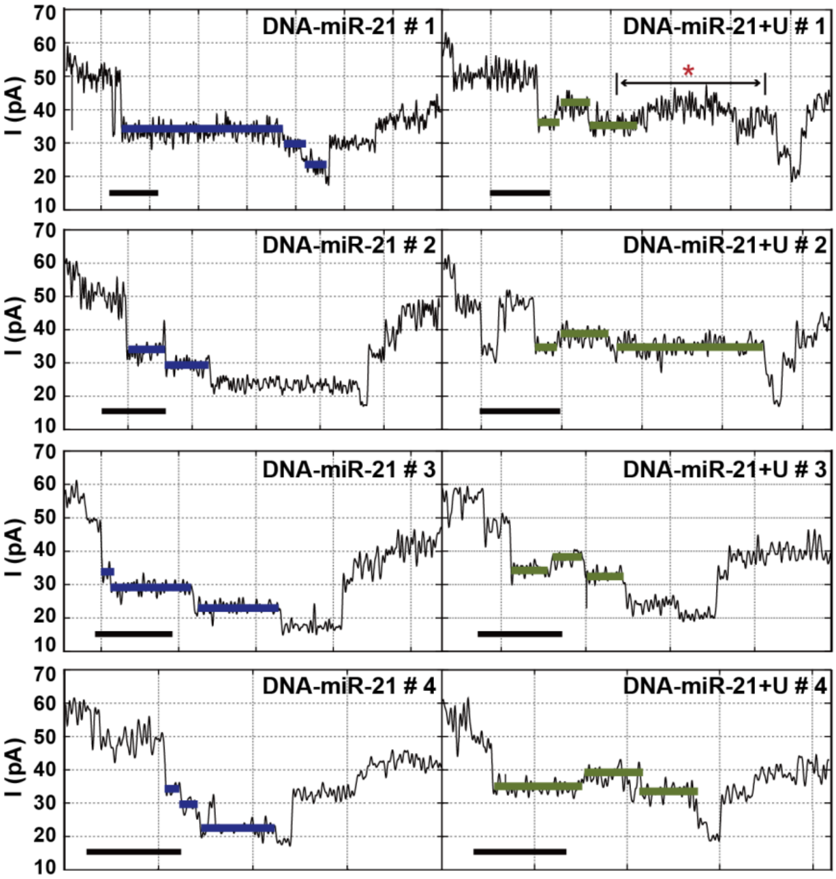

**Fig. S4: Representative current traces for isomiRs discrimination.** Each panel shows individual trace segments, which correspond to the miRNA part of the nanopore sequencing signal of DNA-miR-21 (left) and DNA-miR-21+U (right). Current steps with significant deviations between analytes are indicated by solid lines (blue for miR-21 and yellow for miR-21+U). The "*" indicates polymerase back-stepping[3] that was occasionally observed. The duration time for each nanopore sequencing step is stochastic. Scale bar: 50 ms.

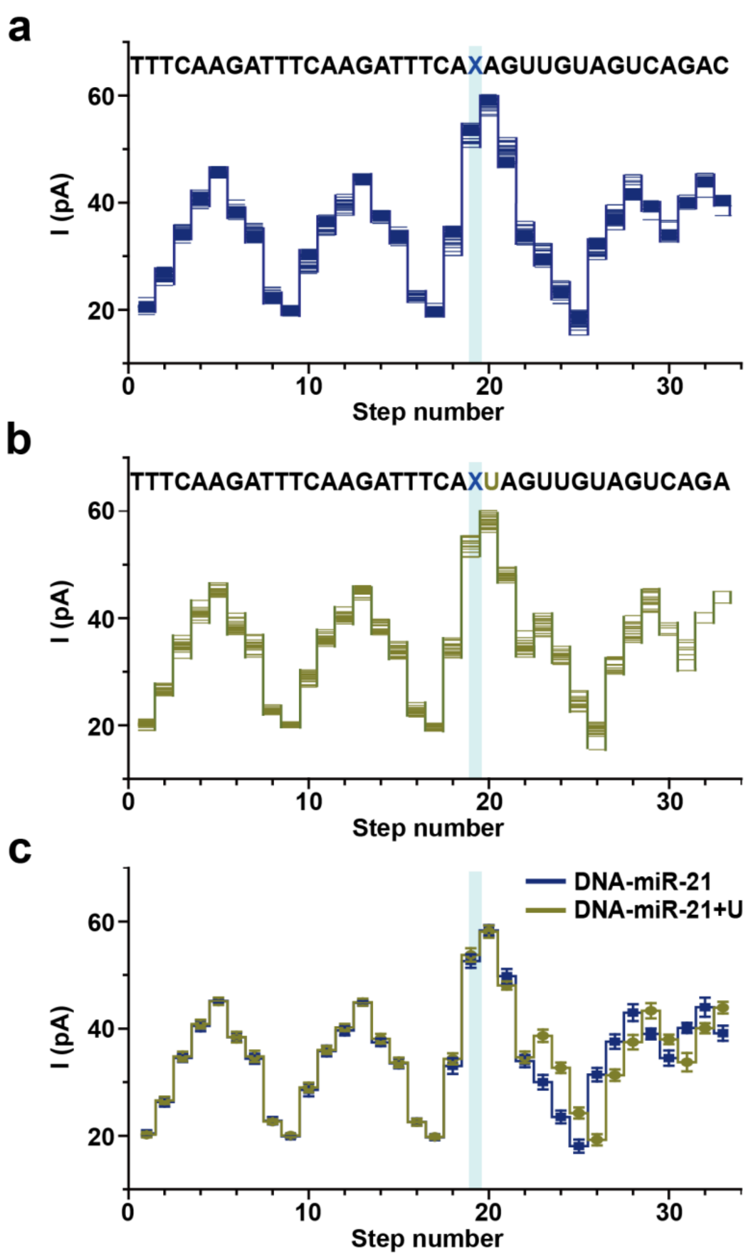

**Fig. S5: Statistical comparison between DNA-miR-21 and DNA-miR-21+U. (a)** Overlay of extracted current steps from multiple DNA-miR-21 events (N=24). **(b)** Overlay of extracted current steps from multiple DNA-miR-21+U events (N=25). The correlated sequences for each sequence are demonstrated on top of the plots in **(a-b)** where an abasic site (X) is located between DNA and miRNA. **(c)** Consensus comparison of sequencing signals between DNA-miR-21 and DNA-miR-21+U. Mean and standard deviations extracted from **a** and **b**. Here, a uridine monophosphate insertion has generated an additional current step when reading DNA-miR-21+U in reference to that of DNA-miR-21. Consequently, the signal pattern from DNA-miR-21+U is shifted by 1 nucleotide. The blue strip in **(a-c)** represents the quadromer reading of TCAX, which is the first quadromer containing the abasic site read by NIPSS.

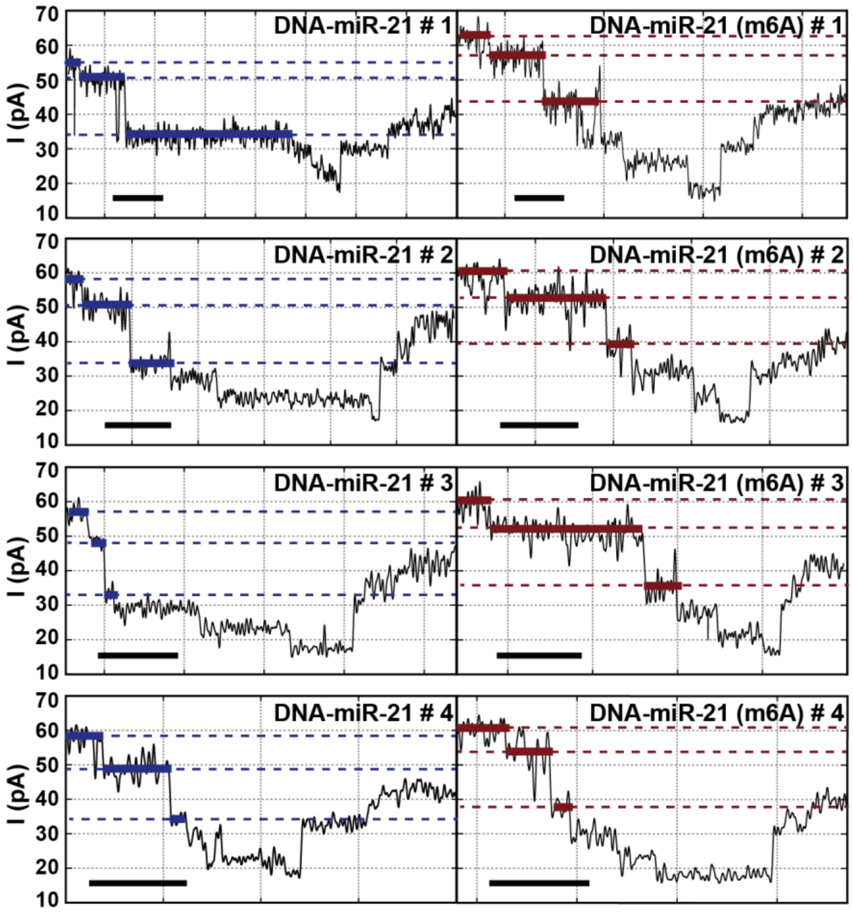

**Fig. S6: Representative current traces for m6A modification detection.** The sequence contexts between DNA-miR-21 and DNA-miR-21(m6A) differ with only one m6A modification **(Table S1)**. Each panel shows individual trace segments, which correspond to the miRNA part of the sequencing signal of DNA-miR-21 (left) and DNA-miR-21(m6A) (right). Current steps with significant deviations between analytes are indicated by solid lines. These level differences could also be recognized from the dashed reference lines. The duration time for each nanopore sequencing step is stochastic. Scale bar: 50 ms.

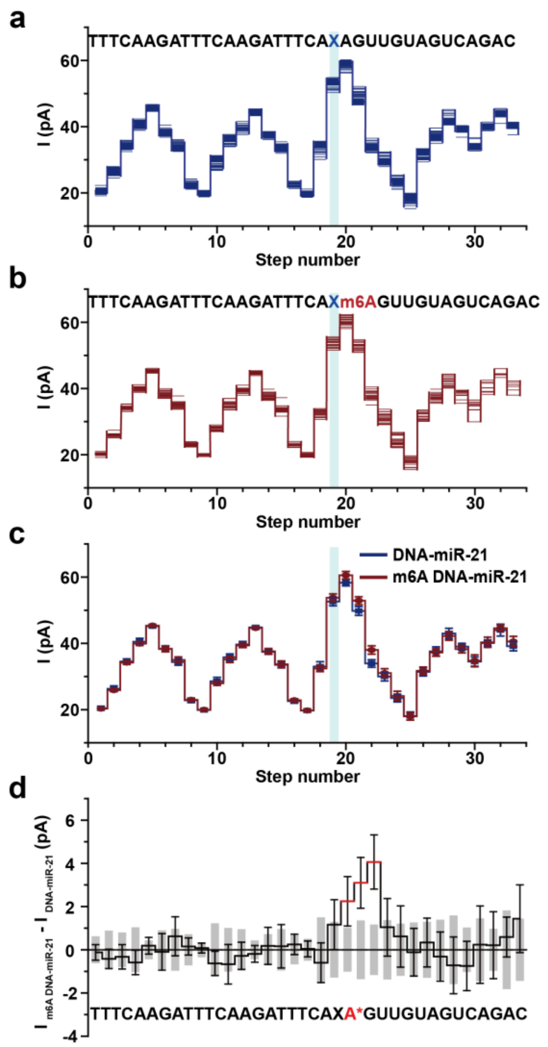

**Fig. S7: Statistical signal variation between DNA-miR-21 and DNA-miR-21(m6A). (a)** Overlay of current steps extracted from multiple DNA-miR-21 events (N=24). **(b)** Overlay of current steps extracted from multiple DNA-miR-21(m6A) events (N=24). Each current step in **(a-b)** represents the mean current values of each quadromer reading. The correlated sequences are shown above the plots. The "X" within the sequence stands for the abasic site located between the DNA and the miRNA segment of the chimeric template. **(c)** Demonstration of different current patterns generated from two DNA-miRNA strands. The average current values and error bars are created from current level traces exhibited in **a** and **b**. The blue strips in **(a-c)** represent the quadromer reading of TCAX, which is the first quadromer containing the abasic site when read by NIPSS. **(d)** Statistical current differences between DNA-miR-21 and DNA-miR-21(m6A). Mean current values in **(c)** were used to construct the current difference map, where the value of $I_{\text{DNA-miR-21(m6A)}} - I_{\text{DNA-miR-21}}$ is displayed by a black solid line. The standard deviation of signals from DNA-miR-21 is shown with gray columns. The standard deviation of signals from DNA-miR-21(m6A) is demonstrated with black bars. Signal variations caused by m6A modification are shown as three red lines. The sequence of the strand is aligned below the figure, where A* (red) represents either A or m6A in the sequence.

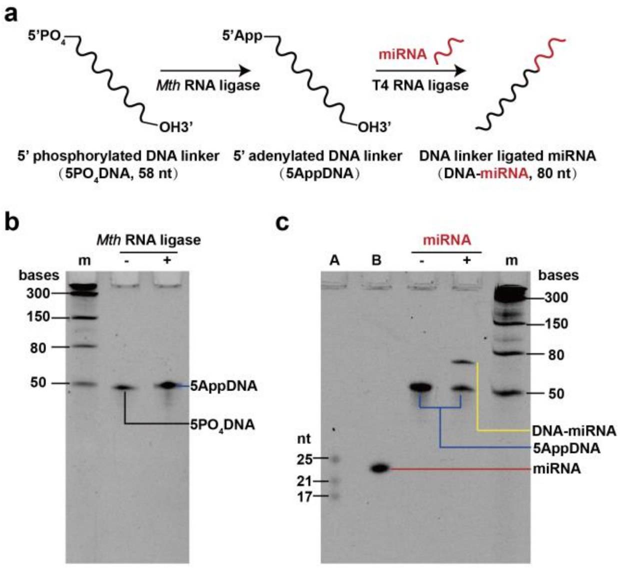

**Fig. S8: Enzymatic ligation between DNA and RNA. (a)** A schematic diagram of enzymatic ligation. The reaction starts with a 58 nt DNA linker with a 5' phosphate group ($5PO_4DNA$). After treatment with the *M*th RNA ligase (New England Biolabs), the 5' end of the DNA linker is adenylated (5AppDNA). To minimize non-specific ligation, T4 RNA ligase 2 truncated K227Q mutant (T4 Rnl2tr, New England Biolabs) was chosen, which specifically ligates the 5' end of the pre-adenylated DNA linker (5AppDNA) with the 3' end of target miRNA. **(b)** Characterization of DNA adenylation by 15% polyacrylamide (PAGE)-urea gel. The adenylation reaction was performed by mixing the following components: 200 pmol $5PO_4DNA$, 4 μL 10x 5' DNA adenylation buffer, 4 μL 1 mM ATP, 4 μL *M*th RNA liagse and nuclease-free $H_2O$ to a final volume of 40 μL. The reaction was incubated at 65 ℃ for 1 h and heat inactivated by incubation at 85 ℃ for 5 min. The reaction product 5AppDNA was purified by ethanol precipitation. Lane + and lane – stand for samples that were incubated with *M*th RNA ligase or not, respectively. Lane m stands for the Low Range ssRNA Ladder (New England Biolabs). 5' adenylation results in slight upshift of the band during gel electrophoresis. **(c)** Characterization of DNA-miRNA ligation. Enzymatic ligation was performed between DNA linker and miR-21+U **(Table S1)**. The ligation results were characterized by 15% PAGE-Urea gel electrophoresis. The ligation reaction was performed by mixing the following components: 5.0 μL 50% (w/v) PEG 8000, 2 μL 10x RNA ligase buffer, 20 pmol 5AppDNA, 10 pmol miRNA, 1 μL T4 Rnl2tr, nuclease-free water to a final volume of 20 μL. The reaction was incubated at 4 °C for 24 h and heat inactivated by incubation at 65 °C for 20 min. Lane A indicates the microRNA Marker (New England Biolabs). Lane B stands for the miR-21+U strand. Lane - stands for the 5AppDNA. Lane + stands for the ligation product. Lane m stands for the Low Range ssRNA Ladder (New England Biolabs). An extra, high molecular weight band was detected in lane +, which is the ligated DNA-miRNA chimeric strand.

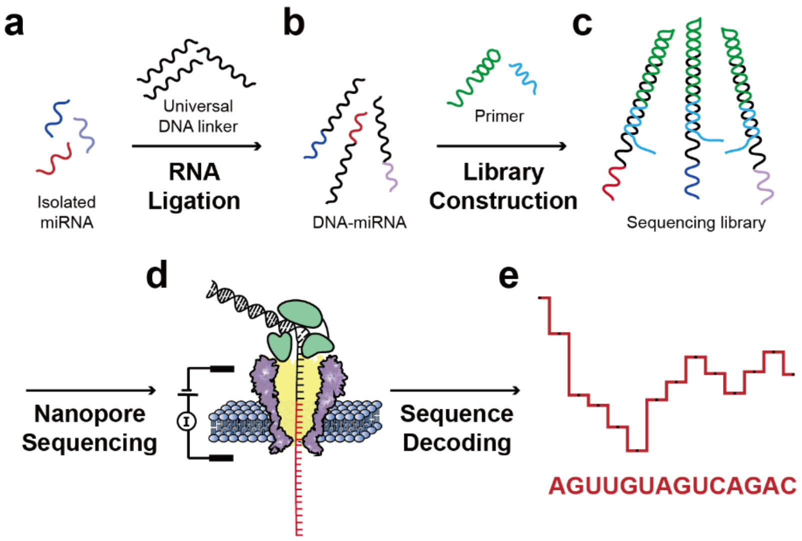

**Fig. S9: A proposed strategy to directly sequence miRNA from natural resources. (a-c)** Schematic diagram of NIPSS sequencing library preparation with isolated miRNA from natural resources. Isolated miRNAs **(a)** could be ligated to form DNA-miRNA chimeric strands **(b)** and subsequently form sequencing libraries **(c)** by thermal annealing. **(d)** Direct miRNA sequencing is carried out as described in this paper. **(e)** Current step transitions during NIPSS reading of miRNA could be decoded into RNA sequences for downstream clinical diagnosis or bioinformatics investigations.

**Table S1: Nucleic acid sequences used in this study.**

| Construct | Sequence (5’-3’) |
|---|---|
| **DNA-miR-21** | UAGCUUAUCAGACUGAUGUUGAXACTTTAGAACTTTAGAACTTTTCAGATCTCACTATCGCATTCTCATGCAGGTCGTAGC |
| **DNA-let-7a** | UGAGGUAGUAGGUUGUAUAGUUXACTTTAGAACTTTAGAACTTTTCAGATCTCACTATCGCATTCTCATGCAGGTCGTAGC |
| **DNA-miR-21+U** | UAGCUUAUCAGACUGAUGUUGAUXACTTTAGAACTTTAGAACTTTTCAGATCTCACTATCGCATTCTCATGCAGGTCGTAGC |
| **DNA-miR-21(m6A)** | UAGCUUAUCAGACUGAUGUUGm6AXACTTTAGAACTTTAGAACTTTTCAGATCTCACTATCGCATTCTCATGCAGGTCGTAGC |
| **Primer** | GCGTACGCCTACGGTTTTCCGTAGGCGTACGCGCTACGACCTGCATGAGAATGC |
| **Blocker** | GATAGTGAGATCTGATTTCCCAAATTTAAA |
| **DNA linker** | $PO_4$-ACTTTAGAACTTTAGAACTTTTCAGATCTCACTATCGCATTCTCATGCAGGTCGTAG-dideoxyC |
| **miR-21+U** | UAGCUUAUCAGACUGAUGUUGAU |

Notes:

1. All miRNA segments of the sequence contexts are underlined.
2. Primer/Template duplex regions in the chimeric template strand are indicated by blue letters.
3. Blocker/Template duplex regions in the chimeric template strand are indicated by red letters.
4. The “X” letter represents the abasic site.
5. $PO_4$ stands for the 5’ phosphate group.
6. dideoxyC stands for a 3’ dideoxycytosine.

**Table S2: Current values of all 4-nucleotides sequence contexts for isomiRs.**

| Level number | miR-21 | | | miR-21+U | | |
|---|---|---|---|---|---|---|
| | Sequence quadromers (3'-5') | Mean I value (pA) | Standard Deviation (pA) | Sequence quadromers (3'-5') | Mean I value (pA) | Standard Deviation (pA) |
| 1 | TTTC | 20.4 | 0.6 | TTTC | 20.2 | 0.5 |
| 2 | TTCA | 26.3 | 0.9 | TTCA | 26.6 | 0.7 |
| 3 | TCAA | 34.5 | 0.8 | TCAA | 34.8 | 0.9 |
| 4 | CAAG | 40.5 | 1.0 | CAAG | 40.9 | 0.8 |
| 5 | AAGA | 45.2 | 0.4 | AAGA | 45.1 | 0.7 |
| 6 | AGAT | 38.4 | 0.9 | AGAT | 38.4 | 1.0 |
| 7 | GATT | 34.4 | 0.9 | GATT | 34.8 | 1.0 |
| 8 | ATTT | 22.8 | 0.7 | ATTT | 22.6 | 0.5 |
| 9 | TTTC | 19.9 | 0.3 | TTTC | 20.0 | 0.3 |
| 10 | TTCA | 28.6 | 1.2 | TTCA | 29.0 | 0.9 |
| 11 | TCAA | 35.8 | 1.0 | TCAA | 36.0 | 0.7 |
| 12 | CAAG | 39.7 | 0.9 | CAAG | 40.2 | 0.7 |
| 13 | AAGA | 44.8 | 0.4 | AAGA | 44.9 | 0.7 |
| 14 | AGAT | 37.4 | 0.7 | AGAT | 38.1 | 0.9 |
| 15 | GATT | 33.5 | 0.9 | GATT | 33.7 | 0.9 |
| 16 | ATTT | 22.6 | 0.6 | ATTT | 22.5 | 0.7 |
| 17 | TTTC | 19.7 | 0.5 | TTTC | 19.8 | 0.4 |
| 18 | TTCA | 33.0 | 1.5 | TTCA | 34.4 | 1.0 |
| 19 | TCAX | 52.6 | 1.3 | TCAX | 53.7 | 1.3 |
| 20 | CAXA | 58.3 | 1.0 | CAXU | 58.1 | 1.1 |
| 21 | AXAG | 49.8 | 1.3 | AXUA | 48.0 | 0.8 |
| 22 | XAGU | 33.9 | 1.1 | XUAG | 34.7 | 1.1 |
| 23 | AGUU | 30.0 | 1.3 | UAGU | 38.7 | 1.2 |
| 24 | GUUG | 23.5 | 1.2 | AGUU | 32.7 | 1.0 |
| 25 | UUGU | 18.1 | 1.2 | GUUG | 24.2 | 1.0 |
| 26 | UGUA | 31.4 | 1.3 | UUGU | 19.2 | 1.0 |
| 27 | GUAG | 37.6 | 1.4 | UGUA | 31.3 | 1.1 |
| 28 | UAGU | 43.0 | 1.6 | GUAG | 37.5 | 1.4 |
| 29 | AGUC | 39.0 | 1.0 | UAGU | 43.3 | 1.4 |
| 30 | GUCA | 34.5 | 1.4 | AGUC | 38.0 | 0.9 |
| 31 | UCAG | 40.1 | 0.9 | GUCA | 33.7 | 1.7 |
| 32 | CAGA | 44.0 | 1.8 | UCAG | 40.1 | 1.0 |
| 33 | AGAC | 39.1 | 1.4 | CAGA | 43.9 | 1.1 |
